# SBIS, a new orange fluorescent vital probe for the 4D imaging of brown algal cells

**DOI:** 10.1101/2025.02.02.636089

**Authors:** Marie Zilliox, Mayeul Collot, Bénédicte Charrier

## Abstract

Living cells of brown algae are difficult to observe in 3D because pigments such as fucoxanthin and chlorophyll diffract light. Furthermore, at the beginning of their life, brown algae develop slowly in seawater. To gain insight into the 3D shape and size of brown algal cells during embryogenesis, we designed a fluorescence probe that efficiently and selectively labels the plasma membrane. Styryl benzoindoleninium sulfonate (SBIS) is a bright orange fluorogenic probe that is soluble and virtually non-emissive in seawater and is activated upon binding to the plasma membrane. Unlike Calcofluor White, SBIS enables observation of cells at thicknesses of up to 25 µm. More importantly, SBIS allows three-dimensional observation of the cells in the growing uniseriate filaments of *Ectocarpus sp.,* the polystichous filaments of *Sphacelaria* rigidula and the cellular monolayered lamina of *Saccharina latissima* over periods of up to seven days. Altogether, these properties allow visualization of the entire cell contours in living brown algae, making the study of early development at the cellular level in 4D now possible in these marine organisms.

## Introduction

Brown algae are multicellular marine organisms whose biology is still relatively unexplored. Nonetheless, they can be found along almost all of the planet’s coasts, where they contribute to local ecology in a number of ways (Arkema et al., 2009; Barbier et al., 2020; Charrier et al., 2012; de Groot et al., 2012; González Fernández et al., 2023; Holdt and Edwards, 2014; Hurd et al., 2022; Krause-Jensen and Duarte, 2016; Malla et al., 2024). Brown algae come in various sizes and shapes (Charrier et al., 2012), ranging from long-bladed kelps measuring over 30 m in length to the microscopic filaments of Ectocarpales species. At first glance, their bodies are made up of a mass of cells of fairly uniform shape and color, and seem poorly differentiated (Charrier et al., 2012; Denoeud et al., 2024). However, looking more closely reveals that their shapes change as they mature: the cuboid epidermal cells of the meristoderm of *Saccharina* (Laminariales) differentiate into spherical and cylindrical cortical cells (Theodorou and Charrier, 2023); the oblong apical and cylindrical subapical cells in the filamentous alga *Ectocarpus* differentiate into spherical cells in the center in the filament (Jia et al., 2017; Le Bail et al., 2008; Le Bail et al., 2011). Even more impressive is the differentiation of the spherical cortical cells into highly branched medullary cells in kelps (Fritsch, 1945; Theodorou and Charrier, 2023). In all these cases, changes in cell shape change is accompanied by a thickening of the cell wall and a change in metabolic function: kelp meristoderm cells are highly dividing cells whereas medullary cells transport metabolites throughout the organism; the apical cell of the filamentous brown alga *Ectocarpus* is primarily dedicated to tip growth and the round, more mature cells in the center of the filament are more active in photosynthesis (Le Bail et al., 2008). Therefore, brown algae have several cell types with different physiological and metabolic functions, and also feature different cell growth rates and shapes. Progress in the understanding of brown algal biology, and especially development, requires imaging these cells in 3D over time.

Brown algal cells are surrounded by a cell wall (CW), which prevents them from moving within the tissue and gives them a specific shape. Because brown algae are not genetically transformable, it is not possible to visualize cell contours using a fluorescent tag encoded in the genome and expressed throughout the development of the organism. Therefore, a fluorescent vital probe harboring chemical characteristics compatible with the biological features of brown algae must be used. In this context, chemical biology is a new exciting approach for developing novel tailored fluorescent imaging tools that meet demanding requirements (Tamura and Hamachi, 2022). Indeed, the probe must be suitable for use in seawater, and be stable over several days, because brown algae grow slowly (the *Ectocapus* filament grows 300 times slower than the pollen tube of tobacco (Rabillé et al., 2019)). However, by far the most important criterion is the ability of the probe to image brown algal cells in 3D, despite the presence of pigments (Muñoz-Miranda and Iñiguez-Moreno, 2023). Carotenoid pigments are abundant in the chloroplasts of brown algal cells (from 26 to 60%) where they absorb blue light (excitation wavelength, 452 nm) and emit red light (emission wavelength, 655 nm; (Wang et al., 2005). In this context, Calcofluor White (CFW) efficiently labels the CW (Clerc et al., 2022; Le Bail et al., 2008), but it has several disadvantages: it stains cellulose, which is not abundant in the walls of brown algae (Charrier et al., 2019; Deniaud-Bouët et al., 2017; Popper et al., 2011). In addition, its excitation wavelength corresponds to UV, whose penetration into living tissues is hampered by light scattering and the high radiation energy level results in significant phototoxicity. Many vital probes are available on the market (for recent reviews, see (Qiu et al., 2025; Tong et al., 2025). However, they often miss their targets in brown algae: despite their morphological resemblance to plants, brown algae are phylogenetically very distant from them (Burki et al., 2020; Silberfeld et al., 2010). This specific evolutionary history confers specific features and components to brown algal cells, making vital probes commonly used in land plants to label the cell contours inefficient. For example, propidium iodide stains the cell wall (CW) in land plants and the nucleus in animal cells, but not in brown algae (observation by B. C.).

To observe the contours of brown algal cells in 3D over several days, we developed the fluorescent vital probe SBIS, which labels the plasma membrane. Here, we show how this probe can be used to monitor the different cell shapes, sizes and growth modes in three species of brown algae, *Sphacelaria*, *Saccharina* and *Ectocarpus*, using confocal and light-sheet microscopy.

## Results

### Chemical and optical characteristic of SBIS

Fluorescent labeling of the plasma membrane (PM) has become relatively common in animal as well as in plant cells (Collot et al., 2022). In particular, we have contributed to this field by developing efficient PM fluorescent markers of various colors, such as the MemBright® family (Collot et al., 2019a; Collot et al., 2019b) and others (Collot et al., 2015; Collot et al., 2020). These probes benefit from balanced hydrophobicity thus allowing fast, fluorogenic and efficient PM staining in various types of samples such as plated cells, 3D samples like spheroids (Collot et al., 2019a), animal tissues including acute brain slices (Collot et al., 2019b) and finally plant tissues (Collot et al., 2020). Moreover, these markers can be used in cells and tissues across various microscopy modes including two-photon imaging (Collot et al., 2019b) and super resolution microscopy, e.g. single molecule localization microscopy (SMLM) (Collot et al., 2019b) and stimulated emission depletion (STED) microscopy (Collot et al., 2020). Our strategy relies on functionalizing a fluorescent dye with an amphiphilic and zwitterionic moiety that mimics lipids and efficiently anchors in the cell lipid bilayer (Collot et al., 2019b). However, this strategy cannot be applied to labeling the PM of marine organisms. The high ionic strength of seawater causes the aggregation and precipitation of these lipophilic probes before they reach their target. Moreover, the thick polar CW of brown algae acts as a shield, preventing the amphiphilic probes from accessing the PM. Therefore, developing such a PM probe is challenging.

To tackle this challenge we synthesized a series of orange-emitting styryl dyes with various degrees of lipophilicity. They were chosen for their ease of synthesis and adjustment (Li et al., 2004; Wuskell et al., 2006); several PM probes including the Fei-Mao (FM) dye family (Schote and Seelig, 1998) and their improved version (Collot et al., 2020) are based on this fluorescent scaffold.

We first evaluated their ability to label the PM of brown algae using epifluorescence microscopy. Among these tested dyes, one demonstrated efficient PM labeling in brown algae: styryl benzoindolenium sulfonate, SBIS (Figure 1A).

**Figure 1:**
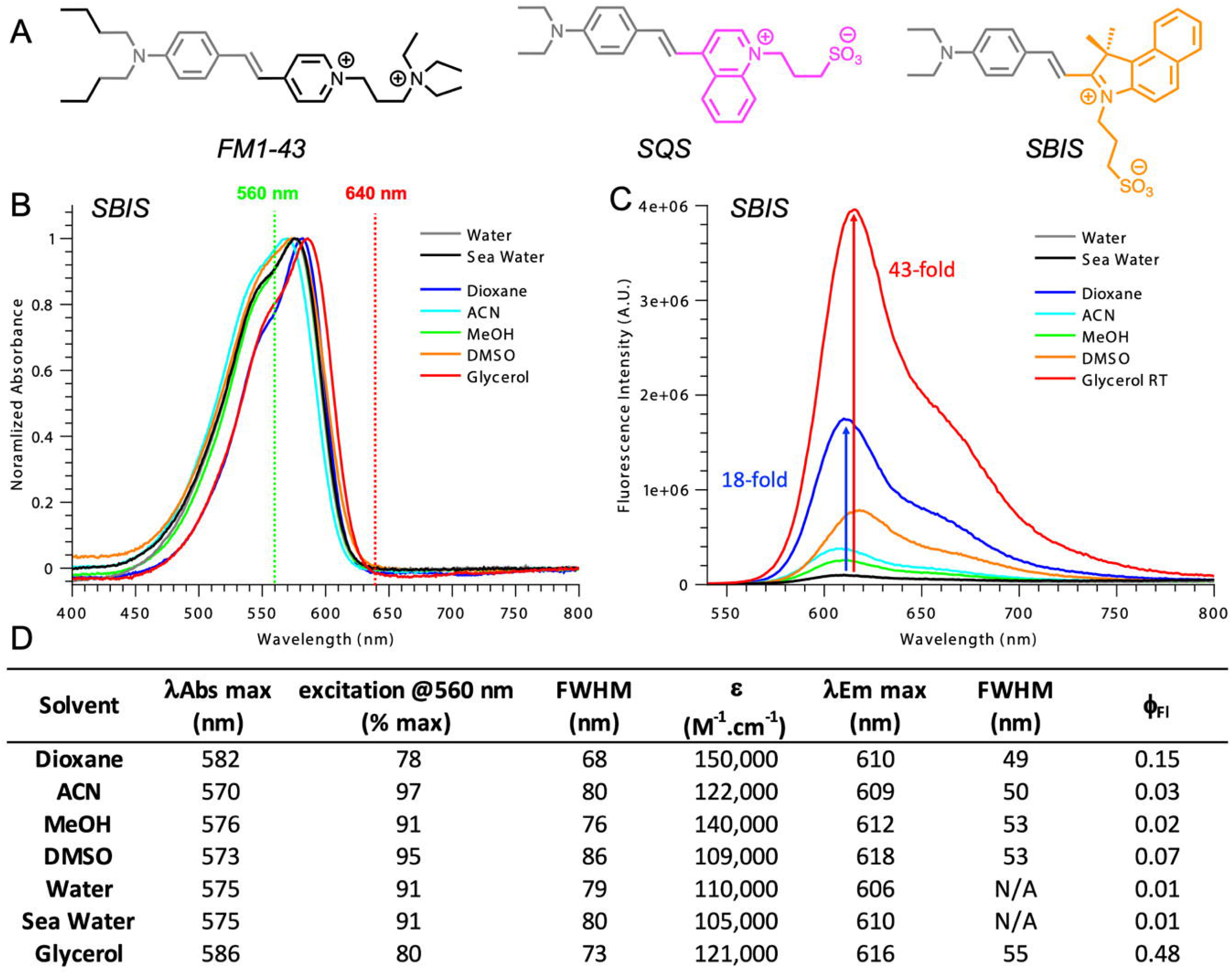
Structural and photophysical properties of SBIS. **(A)** Structures of the styryl dyes discussed in this work. The gray part of the structure is the anillin electron-donor moiety commonly found in the three probes. The quinolinium and benzoindolenium moieties are respectively colored in magenta and orange. The fluorophore acts as the lipophilic group, and the charged moieties act as the polar heads. **(B)** Normalized absorption spectra of SBIS (1 µM) in various solvents. The vertical lines represent the commonly used 560 and 640 nm laser lines. **(C)** Fluorescence intensity of SBIS (1 µM) in various solvents showing the fluorescence enhancement of the dye when placed in a non-polar or viscous environment. Excitation was at 530 nm. **(D)** Photophysical properties of SBIS in the tested solvents. λAbs max and λEm max are respectively the maximum absorption and emission wavelengths. ε is the molar extinction coefficient at the maximum absorption wavelength (λAbs max). FWHM is the full width at half maximum denoting the width of the spectrum. Φ_Fl_ is the quantum yield of fluorescence.

Comparing the structure of SBIS to the other tested dyes revealed that neutralizing the positive charge of the styryl dye with a sulfonate group significantly improved PM labeling ability. Unlike the classical linear structure of a lipid mimic composed of a lipophilic tail and a polar head (like FM dyes; Figure 1A, left), SBIS has a “T” shape with a lipophilic dye and a charged polar head positioned orthogonally (Figure 1A, right). Interestingly, when the benzoindolenium moiety was replaced with a quinolinium moiety, the resulting linear amphiphilic probe SQS (Figure 1A, center) failed to stain the PM, suggesting that the orientation of the probe in the bilayer is a key parameter for effective PM targeting. These observations are in line with a recent study reporting that, for plant cell imaging, T-shaped fluorophores are more efficient than linear ones for staining the PM (Peng et al., 2025). The most likely hypothesis is that the geometry of the probe imposes a more or less pronounced dipole moment, which may lead to significant differences in polarity, and therefore solubility in water, as well as in affinity and orientation in the lipid bilayer. However, further investigations would be necessary, requiring a specific dedicated study.

Following these initial investigations, the photophysical properties of SBIS were characterized in various environments and their absorption and emission spectra were recorded (Figure 1B & C). Compared to other styryl dyes, SBIS displayed unique properties. First, SBIS possesses much higher molar extinction coefficients (ε) than similar styryl dyes (with an aniline donor moiety) like FM1-43 or other styryl dyes we previously developed (Collot et al., 2020). In particular, although those dyes possess an ε around 50,000 M^-1^.cm^-1^, that of SBIS ranged between 105,000 (in seawater) and 150,000 M^-1^.cm^-1^ in dioxane (Figure 1D). Additionally, SBIS exhibited sharp emission peaks with full width at half maximum (FWHM) ranging from 49 to 55 nm, whereas classical styryl dyes exhibit broad excitation bands with FWHM ranging from 80 to 100 nm. Finally, SBIS has a small Stokes shift as low as 28 nm (in dioxane), but styryl dyes are usually characterized by their large Stokes shifts (>100 nm), complicating their use in multicolor imaging. The absorption spectra of SBIS in seawater indicate that SBIS is not prone to aggregation which could be detrimental in imaging.

Interestingly, although SBIS is virtually non-emissive in aqueous media (φFl = 0.01), a fluorescence enhancement of 18-fold was observed in non-polar dioxane (Figure 1C), accompanied by a quantum yield of fluorescence (φFl) of 0.15. In glycerol, a highly viscous polar solvent, SBIS displayed a 43-fold fluorescence enhancement compared with water (Figure 1C) and reached a quantum yield of fluorescence of 0.48.

Importantly, although SBIS is highly excitable at 560 nm, it is virtually non-excitable at 640 nm (Figure 1B). These wavelengths correspond to two commonly used laser lines in fluorescence bioimaging. Hence, SBIS holds promise for use in multicolor imaging with a far-red fluorescent probe.

Overall, SBIS is an orange-emitting dye with outstanding properties similar to those of rhodamine or cyanine dyes rather than styryl dyes, making it highly suitable for bioimaging. Although SBIS is poorly emissive in seawater, it displays high fluorescence enhancements in non-polar and viscous media, which are important features for the efficient staining of the PM of brown algae and for achieving high signal-to-noise imaging.

### SBIS specifically labels the plasma membrane of living *Ectocarpus* sp. cells

Fluorescent probes have become powerful tools to visualize specific cell structures *in vivo*. In brown algal cells, the CW and the PM are located next to each other. To guarantee the specificity of SBIS labeling, we conducted a plasmolysis experiment to detach the PM from the CW in living *Ectocarpus* cells labeled with SBIS and CFW. Plasmolysis revealed that SBIS specifically labeled the PM, compared with CFW which labeled the CW (Figure 2A). In addition, co-labeling of *Ectocarpus* with CFW, SBIS and a commercially used PM fluorochrome FM1-43 show co-localization of SBIS and FM1-43 after plasmolysis, thus ensuring the specificity of SBIS to the PM. Comparison of the signal-to-background ratios of SBIS fluorescence at the PM with that at the cytoplasm provide evidence supporting the qualitative observations indicating that SBIS labels the PM properly (Figure 2B). SBIS also appears to stain fine structures in the PM: in *Ectocarpus*, we observed labeled plasmodesmata, which are cytoplasmic bridges between two cells, and potential Hectian fibrous structures (Yoneda et al., 2020) (Figure 2C).

**Figure 2:**
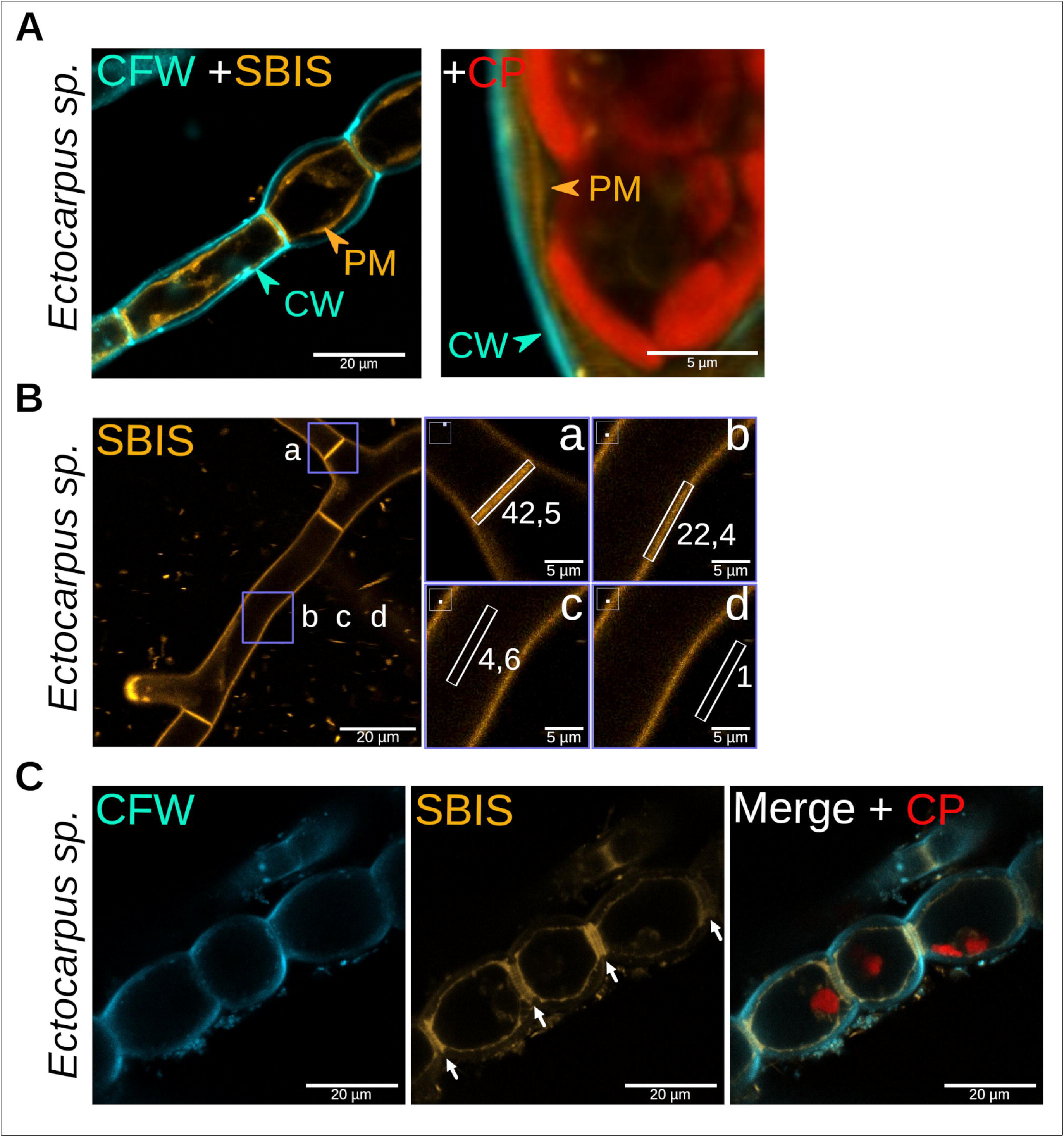
Labeling of the plasma membrane with SBIS. (A) SBIS labels the plasma membrane (PM) of the brown alga *Ectocarpus sp*. Plasmolysis conducted with 2 M NaCl results in the detachment of the PM (orange arrowhead) in response to hyperosmotic stress and reveals the specific labeling of SBIS (orange) on the PM, compared with the labeling of Calcofluor White (CFW, cyan) on the cell wall (CW, cyan arrowhead). (Left panel) Elongated and round cells in *Ectocarpus* filament. (Right panel) Enlargement of an area where the PM has detached. CP: autofluorescence of the chloroplast (red). **(B)** Signal-to-background ratios of the SBIS signal.The intensity of the SBIS signal was measured using Fiji (Schindelin et al., 2012) in areas of the same size (white rectangles) corresponding to **(a)** the PM at a cross-wall, **(b)** the PM along the cell, **(c)** the cytoplasm and **(d)** the external medium (sea water). The ratio of the signal intensity divided by the mean signal found in the background **(d)**, is indicated in the figure. n = 17 samples. **(C)** SBIS allows the labeling of fine structures on the PM, such as plasmodesmata or Hectian fibrous structures (indicated by white arrows). Scale bars are displayed.

*In vivo* live imaging of developing organisms requires that the cells remain alive during the imaging process. However, labeling with fluorochromes may be invasive to the organism. To test the effect of SBIS on brown algal growth and development, we used *Ectocarpus,* which has a growth rate of about one cell division per day (Nehr et al., 2011) and 2.5 µm.h ^-1^ of linear growth at the tip (Rabillé et al., 2019). After labeling *Ectocarpus* with CFW and washing the fluorochrome, *Ectocarpus* was left to grow for five days in the presence of SBIS. After five days, newly formed cells (lacking CFW labeling) cultured in seawater containing only SBIS showed an SBIS-labeled PM and red auto-fluorescent chloroplasts, attesting to their good condition (Figure 3A). Interestingly, when SBIS was washed out of the culture medium before cultivation for five days, we still observed a weak signal corresponding to SBIS in the newly formed cells (Figure 3B). The division rate of SBIS-labeled *Ectocarpus* was measured to be 2.3 µm.h^-1^ (Figure 3A) and 1.55 µm.h^-1^ (Figure 3B). Therefore SBIS does not seem to interfere drastically with *Ectocarpus* growth. The persistence of the signal indicates that SBIS may freely diffuse within the membrane and stain the PM of newly formed cells, or may be recycled in the PM. Alternatively, we cannot exclude the possibility that an exceedingly small amount of SBIS remaining in the medium is enough to label the PM. To confirm the non-toxicity of SBIS in the embryonic development of brown algae and address the possibility that staining delays natural cellular processes, we monitored the development of *Fucus serratus* in the presence of SBIS. Our observation showed that, in the presence of SBIS, the rhizoid of *Fucus* continues to grow and the cells of the embryo divide. In addition, the cell division pattern at 48 h post-fertilization (hpf), thus halfway through the monitoring period, matched that of the *Fucus* plant that has been labeled only with CFW and left to grow before being imaged under a two-photon microscope (Figure 3C).

**Figure 3:**
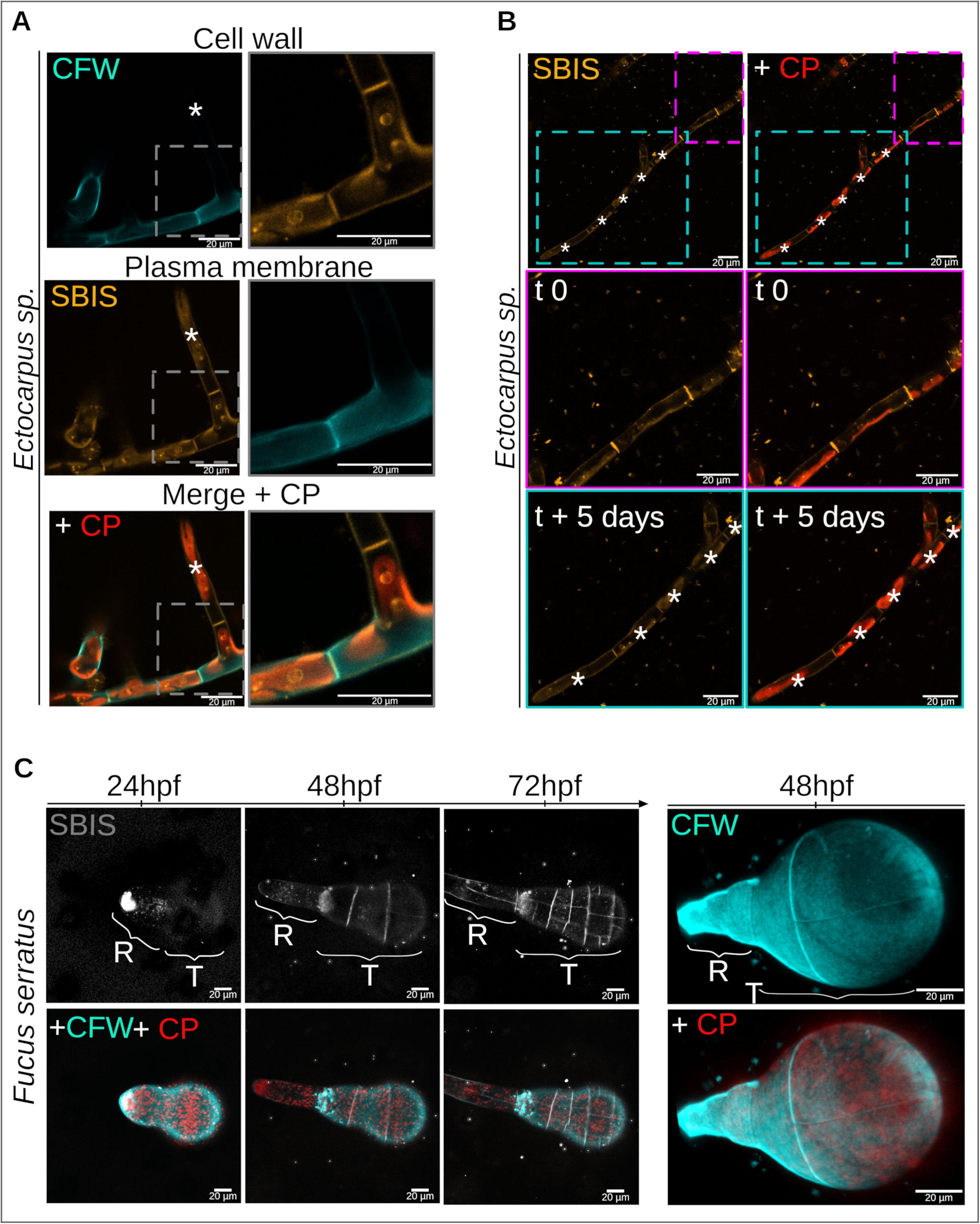
SBIS labeling is non toxic and does not interfere with the growth rate. **(A)** SBIS does not impair the growth of *Ectocarpus sp.*. *Ectocarpus* filaments were labeled with CFW, washed and incubated in SBIS for 5 days. Left: overview of the *Ectocarpus* filament. A branch formed after labeling with CFW and during incubation in SBIS (white asterisk). The branch is labeled with SBIS only. Right: enlargement of the gray-dashed rectangle showing the longitudinal and transverse plasma membrane (PM) of newly formed cells labeled with SBIS. Growth rate is of 2.3 µm.h^-1^. Note the autofluorescent chloroplasts attesting that the cells are photosynthetically active after several hours of incubation in SBIS. The chromatic aberration was corrected in the overlay (merge). **(B)** SBIS accumulates in the PM 5 days after washing. *Ectocarpus* filaments were labeled with SBIS, washed at t0 and grown for 5 days (t+5). Newly formed cells are indicated by a white asterisk. Left: only SBIS signal (orange), Right: SBIS signal and chloroplast auto-fluorescence (CP, in red). Top: overview of an *Ectocarpus* filament. Middle: *Ectocarpus* filament region already present and labeled at t0 (magenta-dashed square). Bottom: New *Ectocarpus* filament cultured for 5 days after removal of SBIS (cyan-dashed square). The five newly formed cells are indicated by a white asterisk. The autofluorescence of the chloroplasts (in red) is an indicator that the cells are in good condition. **(C)** *Fucus serratus* labeled with SBIS and monitored throughout its development. Left: *Fucus* labeled with SBIS and CFW at 24hpf (hours post-fertilization) then washed and monitored for 2 days. The rhizoid (R) grew and the cells in the thallus region (T) divided throughout the time-lapse experiment (only three time points are shown). Right: *Fucus* labeled with CFW at 48hpf, washed out and imaged in two-photon microscopy to visualize the cell wall. CFW is excited with a 405 nm laser and the detection band is 410 –470 nm (Exc 405/Em 410– 470). SBIS: Exc 561/Em 578–632; Chloroplast auto-fluorescence: Exc 561/Em 674–721. In two-photon microscopy, CFW is excited by a 780 nm laser and chloroplasts are excited with a 1080 nm laser. Scale bars represent 20 µm.

Therefore, SBIS can label the PM and be used for monitoring *Ectocarpus* and *Fucus* development *in vivo* and over time.

### SBIS labeling of the brown algal plasma membrane is stable

In this study, we compared the labeling of known commercial PM fluorochromes, i.e. DID (DiIC18(5) solid (1,1’-dioctadecyl-3,3,3’,3’-tetramethylindodicarbocyanine, 4-chlorobenzenesulfonate salt), FM1-43 (N-(3-triethylammoniumpropyl)-4-(4-(dibutylamino) styryl) pyridinium dibromide)) and FM4-64 ((N-(3-triethylammoniumpropyl)-4-(6-(4-(diethylamino) phenyl) hexatrienyl) pyridinium dibromide), with SBIS in three brown algae: *Ectocarpus* sp., *Saccharina latissima* (Figure 4C,D) and *Sphacelaria rigidula* (Figure. 4E,F) We used CFW as a reference to visualize the CW. DID is a far-red fluorescent lipophilic carbocyanine with strong fluorescence when incorporated into membranes. In contrast to SBIS, DID did not label the PM of *Ectocarpus, Saccharina* or *Sphacelaria* cells (Figure 4A,C,E respectively) and co-localized with the auto-fluorescent signal emitted from the chloroplasts (far-red emission signal). We also tested FM styryl dyes, which are widely used for imaging PM and early processes of PM internalization characterized by endocytosis. In contrast to SBIS, FM4-64 labeled the PM, but became internalized almost immediately in *Ectocarpus* and within a few minutes in *Saccharina* and *Sphacelaria* (Figure 4A,C,E, respectively). Similarly, the styryl dye FM1-43 labeled the PM and was quickly internalized by the cells of the cells of the three algae (Figure 4A,C,E, respectively). Altogether, our results show that SBIS labels the PM of brown algae cells and, unlike DID, FM4-64 and FM1-43, is not internalized during acquisition (Figure 4A,C,E). Furthermore, we showed that SBIS remains in the PM for at least 8 days, during which the alga remained alive, as attested by the emission of the expected red light from chloroplasts (Figure 4B,D,F).

**Figure 4:**
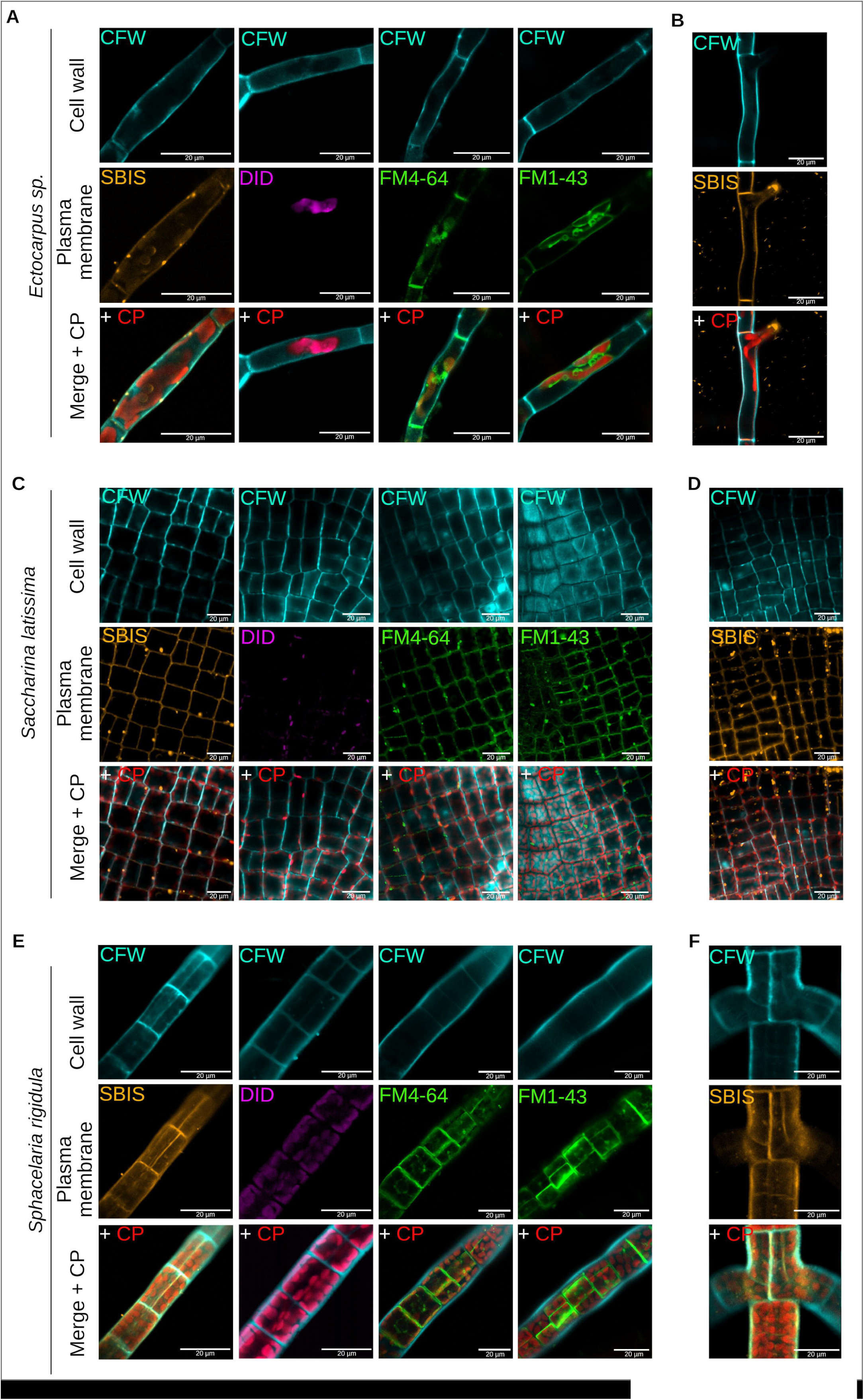
Stability of SBIS in the plasma membrane. Unlike other known plasma membrane (PM) labeling fluorochromes, SBIS is not internalized over time. DID, FM1-43 and FM6-64, were tested in addition to SBIS in three different brown algae: (top) *Ectocarpus sp.*, (middle) *Saccharina latissima* and (bottom) *Sphacelaria rigidula*. **(A)** The PM of the *Ectocarpus sp.* cell is labeled with SBIS, but not with DID. It should be noted that the excitation and emission wavelength ranges of DID are the same as those of chloroplasts, resulting in a co-location of the two signals. FM1-43 and FM6-64 label the PM but are internalized during confocal imaging. **(B)** Long-term retention of SBIS in the PM. SBIS keeps labeling the PM of *Ectocarpus* cells for up to 8 days after SBIS removal by washing. **(C)** In *Saccharina*, the same result is observed as in **(A)**. **(D)** SBIS remains primarily in the PM of *Saccharina* cells for 8 days, but the signal is blurred, suggesting some internalization of the probe. **(E)** In *Sphacelaria*, the same result is observed as in **(A**, **C)**. **(F)** SBIS remains in the PM of *Sphacelaria* cells for up to 8 days. All labeling of PM fluorochromes was done with CFW, which labels the cell wall. Chloroplasts are visualized in red (auto-fluorescence). The chromatic aberrations in the XY plane in the CFW/SBIS overlays (merges) in Figures 4A, E, and F were corrected (see Materials and Methods). The remaining shift between the blue and orange signals is due to the angle of the 2D image extracted from the 3D object reconstruction using Z-stack analysis software. Fig. S1 illustrates this effect. CFW is excited (Exc) with a 405 nm laser and the detection (emission Em) band is 410–470 nm. SBIS: Exc 561/Em 578–632; DID: Exc 633/Em 650–700; FM1-43: Exc 488/Em 580–640; FM4-64: Exc 488/Em 561–641; chloroplast auto-fluorescence: Exc 561/Em 674–721. Scale bars represent 20 µm.

### SBIS labeling combined with confocal, and light-sheet microscopy, enables the observation of the 3D shape of brown algal cells

A major bottleneck to study brown algal cell growth is to observe the shape of these cells in their thickness (third axis). High levels of fucoxanthin pigments are present in the chloroplasts of living algal cells (Haugan and Liaaen-Jensen, 1994), in addition to the phlorotannins present in the cell wall and to a larger extent in the physodes. These are small intracellular vesicles containing secondary metabolites (Ragan, 1976) that are present in the earliest stages of brown algae development (McKinney et al., 2025). These complex and aromatic molecules reflect or scatter light, resulting in poor laser beam penetration into the cells and poor light reflection to the microscope detectors. Hence, we tested the suitability of SBIS for 3D live cell imaging in brown algae, using confocal and light-sheet microscopy. *Ectocarpus* is a brown alga growing only in the X axis, meaning in one dimension, with a cell thickness of 7 µm. Our results show that in confocal microscopy, SBIS allows a clearer visualization of the PM of the cross walls in *Ectocarpus* filaments (Figure 5A,C) and *Saccharina* cells (Figure 5B,D), compared with CFW labeling. Therefore, the use of SBIS to mark the PM in its entirety in the X, Y and Z planes, makes it possible to visualize cell shape in 3D together with internal cell components such as chloroplasts (e.g. Figure 5C,D).

**Figure 5:**
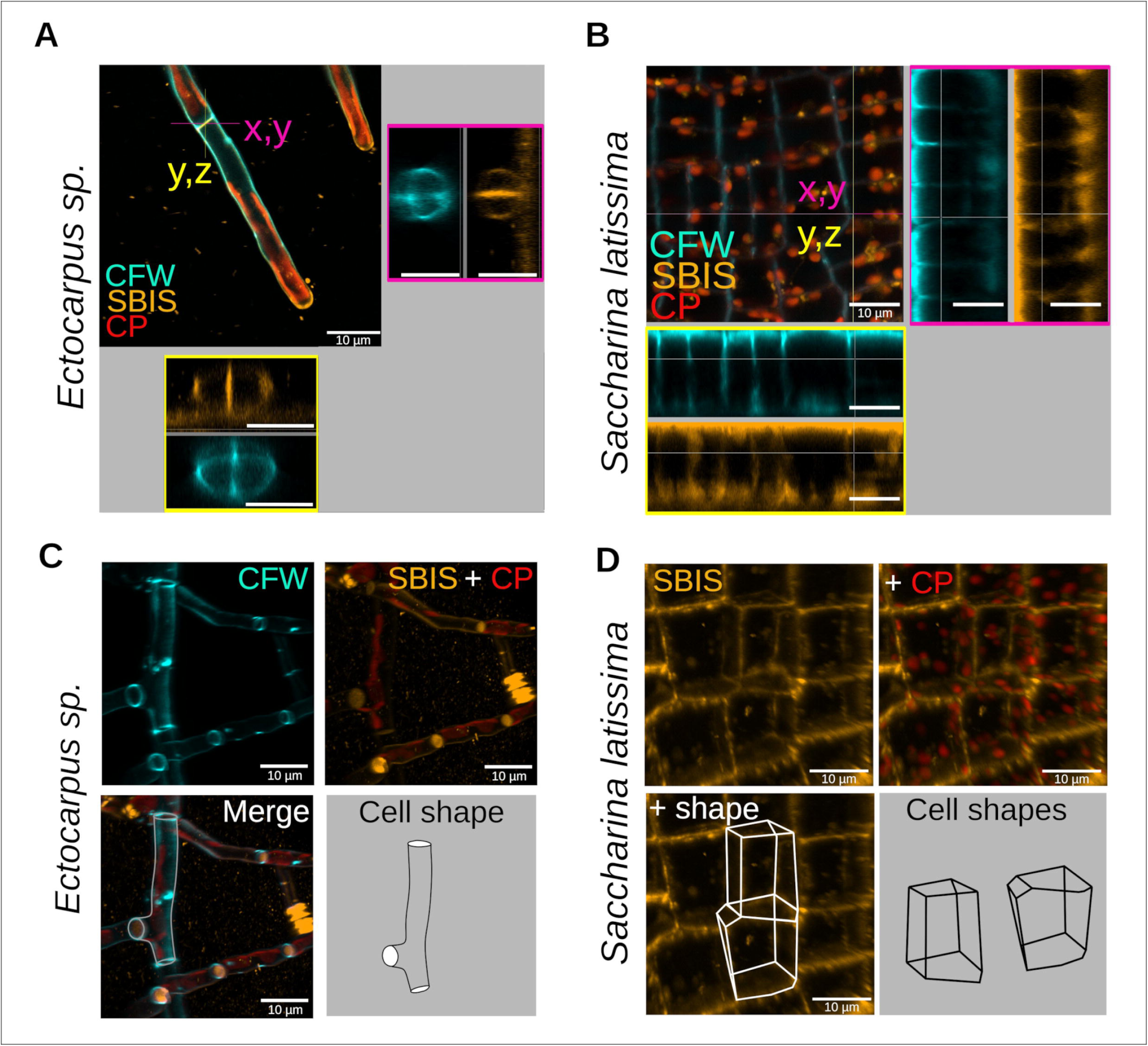
SBIS-mediated depth imaging by confocal microscopy. The ability to image the whole cell PM with SBIS was tested in *Ectocarpus sp.* And *Saccharina latissima*. **(A, B)** Confocal microscopy. The section resulting from orthoslices in the X-Y plane is outlined in magenta, and that in the Y-Z plane in yellow. The orthoslice view seen at the junction of two cells of an *Ectocarpus* filament **(A)** and through a monolayer of cells making the lamina of a *Saccharina* embryo **(B)** shows that SBIS (in orange), in contrast to CFW (in blue) labels the whole cell contour through the thickness of the tissue, which is 7 µm for the *Ectocarpus* filament and 15 µm for the lamina of *Saccharina.* Orthoslices were made with ZEN Blue 3.9 software **(A)** and Imaris 10.2.0 software **(B).** The chromatic aberration was corrected (see Material and Methods). **(C)** Top: 3D reconstruction of a confocal z-stack of *Ectocarpus* labeled with CFW (cyan; left panel) and SBIS with chloroplast autofluorescence (SBIS in orange, CP in red; right panel). Bottom: Merge (left) with cell outlines. 3D reconstruction done using Imaris 10.2.0 software. **(D)** 3D reconstruction of a confocal Z-stack of a region of the *Saccharina* lamina labeled with SBIS (in orange). Chloroplasts (CP in red) are scattered within the cell volume and cell shape is clearly outlined (bottom panels: cell shape schematically drawn). 3D reconstruction done using Imaris 10.2.0 software. CFW is excited with a 405 nm laser and the detection band is 410–470 nm (Exc 405/Em 410–470). SBIS: Exc 561/Em 578–632; chloroplast auto-fluorescence: Exc 561/Em 674–721. Scale bars represent 10 µm.

In the brown algae *Saccharina* and *Sphacelaria*, growth in thickness (Z-axis) takes place before relatively long plateau periods of growth initially restricted in the X or the X, Y axes.

*Saccharina* initiates growth in the Z-axis in a tissue named the transition zone (TZ), which is located at the base of the lamina (Theodorou and Charrier, 2023). It leads to the formation of a meristoderm that expands in thickness during maturation. Due to their location in thick tissue, imaging cells in the TZ is extremely challenging. We mounted *Saccharina* embryos in the light-sheet and focus on imaging the TZ with the acquisition of z-stacks (Figure 6Aa). Orthoslice views in the X-Y axes through the TZ showed that the CFW signal was visible mainly in the outer cell wall (i.e. at the surface of the embryo) and a few µm into the internal, anticlinal cell walls (Figure 6Ab, Z-axis shown in middle panel outlined in blue, X-Y plane in the middle panel outlined in pink). In comparison, SBIS was able to label the PM of the cells in the TZ, allowing us to distinguish the cell shape of at least the outermost layer of cells (Z-stack outlined in blue and orthoslice outlined in pink, showing up to two layers of cells; Figure 6Ab, with LUT rendering helping with viewing on SBIS, right-handed side panel). SBIS clearly enhanced the visualization of cells located deeper in the TZ of the *Saccharina* lamina. *Sphacelaria* also develops a 3D tissue through successive and perpendicular longitudinal cell divisions of the subapical cell located under the apical (Ap) cell. Hence, new transverse membranes located deeper inside the tissue are produced along the filament (schematized in Figure 6Ba). Our results show that SBIS, unlike CFW, was able to label the PM of the first cell dividing in 3D (Figure 6Bb upper panel). Then, 202 µm further down along the filament, SBIS was able to display transverse PM resulting from an additional longitudinal cell divisions (Figure 6Bb bottom panel).

**Figure 6:**
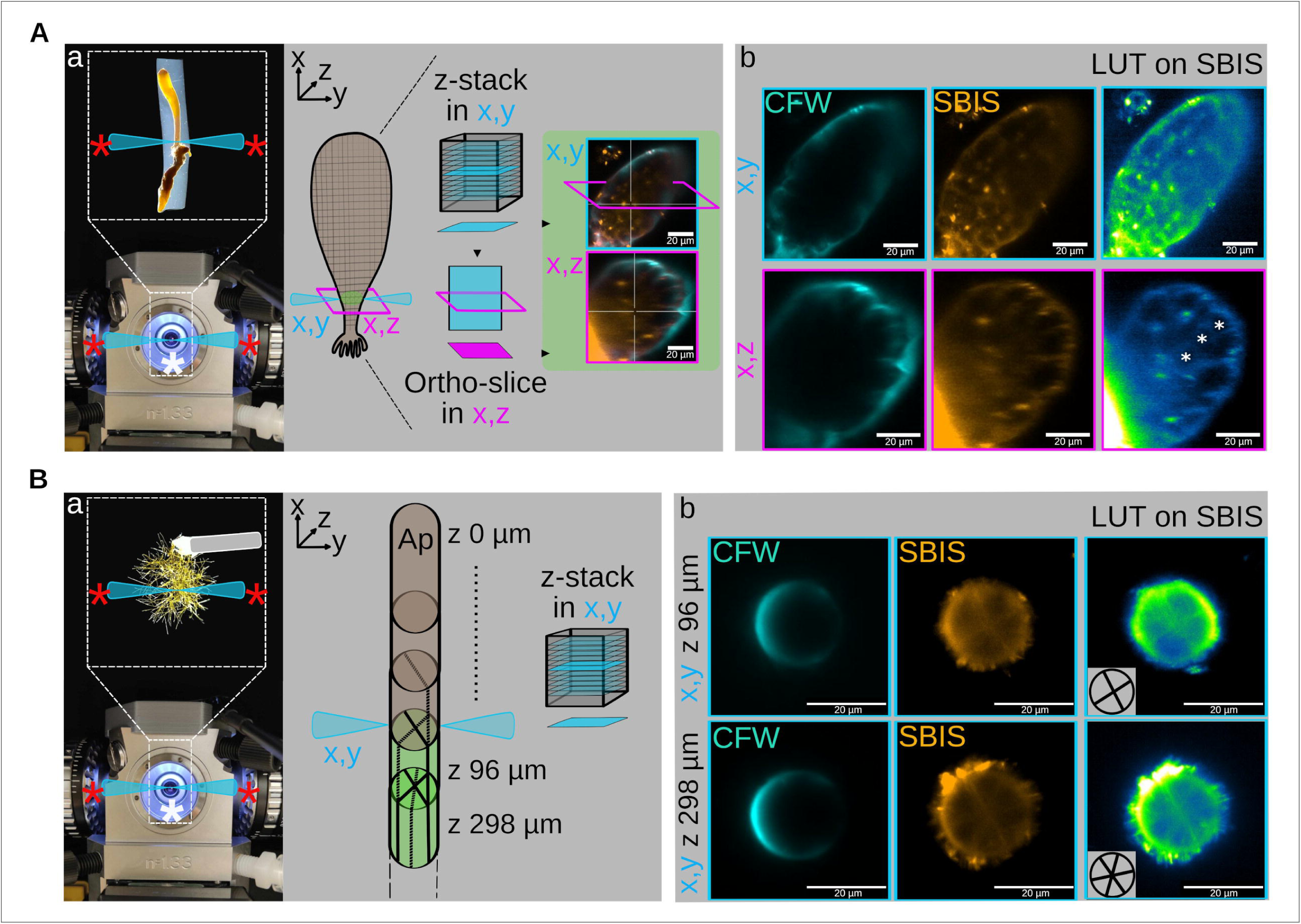
SBIS-mediated depth imaging in light-sheet microscopy. The ability of SBIS to label cell thickness was tested in the brown algae *Saccharina latissima* and *Sphacelaria rigidula*. **(A)** Imaging of a multilayered 20-day old embryo (corresponding to early Phase III as defined in Theodorou and Charrier, 2023) of *Saccharina* in 3D using light-sheet microscopy. **(a)** Mounting of *Saccharina* in agarose between two illumination objectives (red *, light sheets schematized in blue) and in front of an observation objective (white *). Schematics of *Saccharina* with the two light sheets (schematized as blue triangles) placed to image the multilayered transition zone (TZ, green circle). XZ orthoslice (in magenta) was carried out at the middle of the TZ (photo on the right shows the Calcofluor White (CFW) signal in cyan and the SBIS signal in orange). Blue boxes: section in the X/Y plane; Magenta boxes: section in the X/Z plane. **(b)** Top panel: view in X,Y showing the TZ labelled with CFW (cyan) and SBIS (orange) between the light sheets (Z-stack; boxed in cyan). Bottom panel: view in X,Z perpendicular to X,Y (XZ orthoslice boxed in magenta). CFW only labels the outer cell wall of the epidermal cells, whereas SBIS labels several cell layers. A Green Fire Blue LUT inferred from Fiji on the SBIS channel can help distinguish the PM. The three cell layers are displayed by asterisks. **(B)** Imaging of *Sphacelaria* cells in 3D using light-sheet microscopy. **(a)** Mounting of *Sphacelaria* glued on a glass capillary to localize a filament between the two illumination objectives (red *, light sheets schematized in cyan) and in front of an observation objective (white *). Schematics of a *Sphacelaria* filament, cells below the apex (Ap) located at z0 µm develop progressively in 3D (in green). Hence, the light sheets (cyan triangles) are placed paralleled to the filament to image these cells in X,Y (Z 96 µm and Z 298 µm). **(b)** Unlike CFW (cyan), SBIS (orange) labels the transverse PM of the first *Sphacelaria* cell developing in 3D located in Z 96 µm (top panel) and in z298 (bottom panel). The inferred position of the new transverse PMs is displayed in the insert in the bottom left-hand corner of the LUT rendered image. CFW was excited with a 405 nm laser and the detection wavelength is 415 nm (Exc 405/Em 415). SBIS: Exc 561/Em 571 nm; chloroplasts: Exc 561/Em 571 nm. Scale bars represent 20 µm.

Taken together, our results demonstrated that SBIS is a powerful fluorescent probe for *in vivo* 3D imaging of brown algal cell contours in multicellular tissues.

## Discussion

Brown algae are ubiquitous macroscopic marine organisms, yet very little is known about the mechanisms involved in their growth (Charrier et al., 2012). From embryogenesis on, they can develop complex tissues in the form of embryos comparable to those of animals and plants in their primary architecture, as do the algae of the order Fucales (Berger et al., 1994; Brownlee and Berger, 1995). Others grow in successive stages, during which cell growth and division take place in only one orientation, before switching to another spatial dimension, resulting in wider and thicker tissues (Boscq et al., 2024a; Boscq et al., 2024b; Theodorou and Charrier, 2023). In this respect, we recently showed that the cells of the embryonic lamina of the kelp *Saccharina latissima*, in order to initiate growth in thickness, first elongate perpendicularly to the surface of the lamina until they reach ∼30 µm in length, before dividing parallel to this surface (Theodorou and Charrier, 2023). To identify the process that controls the orientation of growth during the embryogenesis of these organisms, it is essential to visualize cell shape in 3D over time, as done in land plant embryos (Laruelle et al., 2022).

However, CFW, which labels the cellulose present in the CW, cannot be used to image living brown algal cells thicker than 7 µm using confocal microscopy. Propidium iodide is often used to delineate the outline of plant cells. However, in brown algal cells, it only stains small, compacted structures (most likely vesicles) (data not shown), whereas in animals, it stains the nucleus.

We have also shown that the FM4-64 and FM1-43 plasma membrane markers internalize quickly and do not allow the cell outline to be imaged for more than a few minutes. Other currently available probes can label the plasma membranes of plant cells. The near-infrared probe APMem-1 based on aggregation-induced emission is one example, but it has been reported to allow imaging for no longer than 10 h (Zuo et al., 2023). In the kelp *Nereocystis luetkeana,* another lipophilic fluorescent dye, DiOC6 (3,3′-dihexyloxacarbocyanine iodide; Molecular Probes, Novato, CA, USA), was shown to label the intracellular membranes (Knoblauch et al., 2016) with excitation and emission spectra (488/505 nm) distinct from CFW and chloroplast auto-fluorescence. However, it was not shown whether DiOC6 was capable of remaining in the PM of living cells for several days or if it was phototoxic (Fotopoulos, 2012).

Here, we identified and characterized SBIS, a new fluorescent probe that stably stains the contours of brown algal cells in 3D using both confocal and light-sheet microscopy. SBIS remains localized in the PM for at least 8 days, making the monitoring of cell growth and division possible during algal development. FM4-64 has been reported to be internalized in several studies in brown algae, especially *Fucus* (e.g. (Belanger and Quatrano, 2000) and *Sphacelaria* (e.g. Aoki et al., 2023) and here we show that FM1-43 internalizes even faster than FM4-64 in brown algal cells. The reasons why SBIS escapes internalization are obscure and, in general, cellular trafficking of fluorescent probes lacks straightforward explanations. The charges and hydrophobicity of the probe likely play an important role. For example, for the same plasma membrane target, uncharged BODIPY-based probes (Collot et al., 2019a) offer much more persistent membrane labeling than positively charged cyanine-based probes (Collot et al., 2019b). In our study, FM1-43 is doubly positively charged, whereas SBIS is zwitterionic and therefore neutral overall. Also, as described above, SBIS has a different geometry from the other probes, which may also affect its internalization.

These differences in charge and spatial conformation may explain why some probes are internalized and others are not. In particular, in other organisms, internalization of extracellular cargos follows two endocytosis pathways, one depending on clathrin proteins (Johnson, 2024) and the other one through the formation of caveolae involving proteins such as caveolin and cavins (Matthaeus and Taraska, 2021). Selective internalization depends on specific membrane nanodomains (Jaillais et al., 2024) that in some cases have been reported to contain specific proteins and lipids such as sphingolipids and cholesterol present on the surface of the plasma membrane (Róg and Vattulainen, 2014). A non-interaction of SBIS with these specific proteins or lipids might explain why SBIS is not internalized. In brown algae, the fast (a few minutes), selective internalization of FM1-43 (and of FM4-64 to a lesser extent) compared with the long-lasting maintenance of SBIS on the cell surface is most likely based on clathrin-dependent mechanisms, because homologs of members of this protein machinery are present in the genome of brown algae, in contrast to caveolins and cavins (BLAST search in BOGAS *Ectocarpus* genomics database (https://bioinformatics.psb.ugent.be/orcae/overview/Ectsi (Cock et al., 2010) and the Phaeoexplorer database for 40 brown algal genomes; https://phaeoexplorer.sb-roscoff.fr/home/ (Denoeud et al., 2024)).

The diversity of endocytotic mechanisms that has been illustrated in animals, plants and yeast (Jaillais et al., 2024; Johnson, 2024; Tsai et al., 2020) opens the door to the possibility that brown algae have developed specific alternative mechanisms, in addition to the clathrin-dependent pathway.

Furthermore, the spectrum of SBIS does not overlap that of chloroplast auto-fluorescence or CFW, which offers the possibility of imaging a third cellular component, i.e., the PM, in addition to the CW and chloroplasts. This aspect is particularly interesting because chloroplasts are often pressed against the PM of brown algal cells and, due to the low Stokes shift of SBIS, they can be distinguished from each without overlapping. SBIS can be thus used in combination with a blue, green and red fluorochrome to obtain multicolor images using separate channels, revealing several structures in the cell.

With the aim of imaging the cellular forms of brown algae over time, SBIS complements the CarboTag marker, which was recently developed to stain the pectins in land plant CWs, and which, surprisingly, stains the pectin-free wall of brown algae (Besten et al., 2025). This example further suggests that the identification of vital fluorescent probes for imaging brown algal cells may require an empirical approach, and testing a wide range of candidate molecules.

SBIS will now enable 4D imaging of cells during the embryogenesis of brown algae, making the growth dynamics and geometry accessible to quantification. These data will feed mechanical models that simulate the formation of 3D tissues in brown algae, which has never been undertaken before. However, mechanical modeling of *Ectocarpus* tip growth (Rabillé et al., 2019) and cell rounding (Jia et al., 2017) has already laid the groundwork. To obtain cell size and shape in 3D over time, the images obtained from *Ectocarpus*, *Saccharina* and *Sphacelaria* will need to be processed by image analysis software. In addition to existing 3D reconstruction software, automated image analysis and segmentation algorithms based on deep learning are being developed specifically for brown algae images.

Finally, given the appealing features and excellent performance of SBIS in complex, pigment-rich samples such as brown algae, and in light of the current lack of fluorescent probes that are compatible with seawater, SBIS may find a broader application in the imaging of other types of marine species. Therefore, it will be made available to the marine biology community, thus expanding on the range of imaging tools in this field.

## Materials and methods

### Preparation of SBIS

#### Materials for probe synthesis

All starting materials for synthesis were purchased from Sigma-Aldrich, BLD Pharm or TCI Europe and used as received unless stated otherwise. All the synthesized compounds were monitored on TLC (silica) and purified using silica gel columns (Interchim Silica HP, 30 µM) on a Puriflash 520X+ (Interchim). NMR spectra were carried out on 400 and 500 MHz Bruker Advance II spectrometers. The ^1^H and ^13^C NMR chemical shifts are reported relative to residual solvent signals. Data are presented as follows: chemical shift (ppm), multiplicity (s = singlet, d = doublet, t = triplet, q = quartet, p = pintet, dd = doublet of doublets, m = multiplet, br = broad), coupling constant *J* (Hz) and integration. Mass spectra were obtained using an Agilent Q-TOF 6520 mass spectrometer and the MALDI-TOF spectra were collected on a Bruker Autoflex II TOF/TOF mass spectrometer (Bruker Daltonics, Billerica, MA, USA).

#### Spectroscopy

Milli-Q water (Millipore) and artificial seawater (marine sea salt: Tropic Marin Classic Meersalz) were used for spectroscopy. All the solvents were spectral grade. Absorption and emission spectra were recorded at 20°C on a Cary 4000-HP spectrophotometer (Varian) and a FluoroMax-4 spectrofluorometer (Horiba Jobin Yvon) equipped with a thermostated cell compartment, respectively. For standard recording of fluorescence spectra, emissions were collected 10 nm after the excitation wavelength. All the spectra were corrected from the wavelength-dependent response of the detector. The fluorescence quantum yields ΦFl were determined from the following equation:

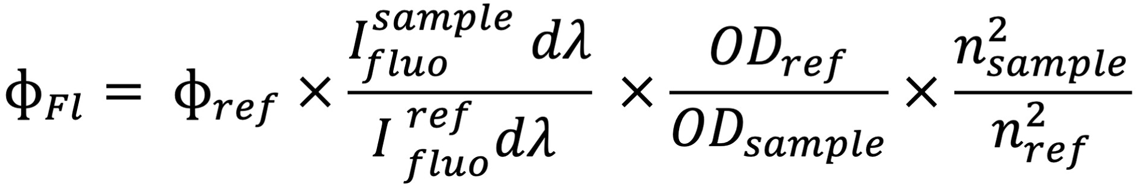

where OD is the optical density at the excitation wavelength, and η the refraction index of the solvent. Rhodamine B was used as the reference (in water, ΦFl= 0.31) (Magde et al., 1999). The oil mixtures were prepared as follows: 1 g of the appropriate oil ratio was heated to 50°C. 5 µL of a 200 µM solution of the dye in dioxane was added and the mixture was homogenized manually. The samples were cooled down at room temperature before being analyzed.

#### Synthesis of SBIS

3-(1,1,2-trimethyl-1*H*-benzo[*e*]indol-3-ium-3-yl)propane-1-sulfonate **1** was synthesized according to a protocol described in the literature (Menéndez et al., 2013).

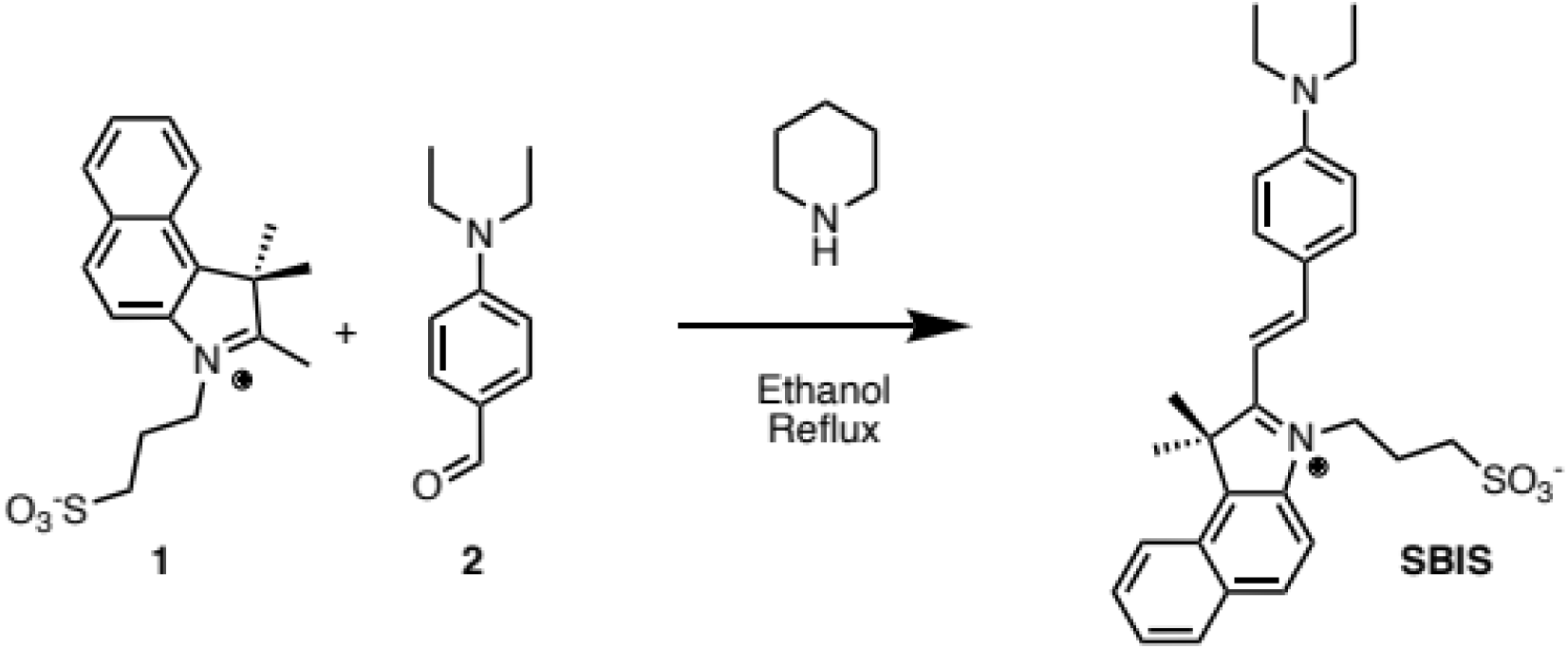

Piperidine (0.5 mL) was added to a solution of 3-(1,1,2-trimethyl-1*H*-benzo[*e*]indol-3-ium-3-yl)propane-1-sulfonate **1** (500 mg, 1.51 mmol) and 4-(diethylamino)benzaldehyde **2** (320 mg, 1.81 mmol, 1.2 eq) in absolute ethanol (30 mL). The solution was refluxed at 110°C and rapidly turned dark violet. The reaction was allowed to reflux for 1 h before being cooled down to room temperature. The solvents were evaporated under reduced pressure and the crude fraction was purified by column chromatography on silica gel (DCM:MeOH, 99:1 to 9:1) to obtain 350 mg of SBIS as a dark violet solid (Yield, 47%). Rf = 0.38 (DCM:MeOH, 9:1).

^1^H-NMR (400 MHz, MeOD): δ 8.40 (d, *J* = 15.5 Hz, 1H, CH ethylene), 8.35 (d, *J* = 8.5 Hz, 1H), 8.16 (d, *J* = 8.9 Hz, 1H), 8.10 (d, *J* = 8.5 Hz, 1H), 8.09-8.05 (m, 2H), 7.92 (d, *J* = 8.9 Hz, 1H), 7.74 (ddd, *J* = 8.4, 7.0, 1.3 Hz, 1H), 7.62 (ddd, *J* = 8.1, 7.0, 1.0 Hz, 1H), 7.45 (d, *J* = 15.5 Hz, 1H, CH ethylene), 6.92 (d, *J* = 9.2 Hz, 2H), 4.81 (t, *J* = 8.2 Hz, 2H, CH2-N^+^), 3.61 (q, *J* = 7.1 Hz, 4H, 2 CH2 Et), 3.10 (t, *J* = 6.3 Hz, 2H, CH2), 2.42 (tt, *J* = 9.6, 5.0 Hz, 2H, CH2), 2.08 (s, 6H, 2 CH3 Me), 1.29 (t, *J* = 7.1 Hz, 6H, 2 CH3 Et).

^13^C-NMR (101 MHz, MeOD): δ 190.62, 180.50, 154.31, 153.61, 138.87, 136.32, 133.09, 130.90, 129.83, 127.74, 127.58, 125.95, 122.50, 122.30, 112.12, 111.53, 103.21, 52.65, 44.70, 43.92, 31.62, 25.90, 22.31, 11.52.

HRMS (ESI^+^), calculated for C_29_H_35_N_2_O_3_S^+^ [M+H]^+^ 491.2363, found 491.2387

### Culture of algae

*Ectocarpus* sp. material was produced from fragmentation of the WT male strain Ec32 (CCAP 1310/4; origin San Juan de Marcona, Peru) parthenosporophytes.

Embryos of *Saccharina latissima* were produced from fertilization of the female gametophyte strain F1 with the male strain M1 as described in (Boscq et al., 2024b). Female gametophytes of *Sphacelaria rigidula* were grown from fragmentation of adult cultures. All alga thalli were grown in full-strength Provasoli-enriched (Starr and Zeikus, 1993) autoclaved natural or artificial seawater (pH 7.8) (Tropic Marin®) in a culture cabinet at 13°C with a 12:12 light: dark cycle (light intensity, 29 µmol photon·m ^-2^·s^-1^) as described in (Le Bail and Charrier, 2013).

Fertile thalli of female and male *Fucus serratus* were obtained from EMBRC-France (Roscoff marine station, Sorbonne University, France), from which embryos were produced as indicated in (Siméon and Hervé, 2017).

### Fluorochromes

Brown algae were labeled with Calcofluor White (ref BCCK5631 from Sigma-Aldrich ®) at 10 µM, SBIS at 5 µM, DID at 10 µM, FM6-64 at 10 µM, FM1-43 at 10 µM, in seawater overnight at 13°C and washed 3 times in seawater and 13°C the next morning. DAPI was used at 20 µM for 1.5 h at 13°C and then washed 3 times in seawater. DID (ref. D7757), FM6-64 (ref. T3166) and FM1-43 (ref. T3163) were purchased from ThermoFisher Scientific®. DAPI (ref. D9542) was purchased from Sigma-Aldrich®. The excitation and emission spectra (λmax) of the different probes are as follows: DAPI: 405/488 nm; CFW: 405/433 nm; SBIS: 561/ 610 nm (Fig. 1); FM1-43: 488/590 nm; FM4-64: 488–514/625–670 nm; DID: 640/668 nm; chlorophyll A (Chloroplasts): 430–561 / 650–700 nm. Staining and observations were carried out at least three times for each dye.

### Confocal microscopy

The algal samples were imaged on a Zeiss LSM 880 confocal microscope with a LD C-Apochromat 40x/1.1 W Corr M27 objective (water as immersion medium). *Ectocarpus*, *Saccharina, Sphacelaria* and *Fucus* were mounted in a 20 mm diameter glass-bottom cell culture dish (NEST Biotech), glass of standard thickness 0.16-0.19mm (Cat. No. 801001) filled with seawater. For every alga, co-labeling of CFW + SBIS or DID or FM4-64 or FM1-43 was performed on the same day and imaged in the same glass-bottom cell culture dish.

A shift due to chromatic aberration was observed between the different emitted wavelengths. Fluorescent beads (Fluorebrite® Multifluorescent Microspheres, 1.00 µm) were used to measure the shift between blue and orange/red light, and all merged images were corrected in Fiji using the ‘Image > Transform > Translate’ function’.

### Light-sheet microscopy

The algal samples were imaged on a Zeiss Lightsheet 7 microscope using a W Plan-Apochromat 20x/1.0 DIC M27 objective. *Saccharina* was mounted in 1% low melting agarose (ref SLCN4676 from Sigma-Aldrich ®) in seawater and *Sphacelaria* was glued (ref. 62928642 from Leroy Merlin ®) onto a glass capillary.

### Imaging: software used for 3D image reconstruction

Z-stacks and snaps taken on the LSM880 and LS7 in their native format (.czi) were visualized in Fiji (open source, available at http://fiji.sc/Fiji; (Schindelin et al., 2012)), Zen Blue (Carl Zeiss Microscopy, LLC) and Imaris 10.2.0 (BitPlane, South Windsor, CT, USA). Quantification of signal intensity was carried out in Fiji.

## Acknowledgements

We thank Tanguy Dufourt for the initial labeling experiments. We thank Jacques Brocard and the PLATIM platform at SFR Biosciences (Universite Claude Bernard Lyon 1, CNRS UAR3444, Inserm US8, ENS de Lyon) for providing access to the Imaris 10.2.0 imaging analysis software. Part of the work was funded by the European Union (ERC, ALTER e-GROW, project number 101055148). Views and opinions expressed are those of the author(s) only and do not necessarily reflect those of the European Union or the European Research Council Executive Agency. Neither the European Union nor the granting authority can be held responsible for them. For purposes of open access dissemination, the authors applied for a CC BY-NC license for this document.

## Contribution

M.C. produced SBIS and M.Z. designed and carried out all the image acquisitions. M.C, M.Z. and B.C. directed the work and wrote the article. All the authors read and approved the manuscript.

## Notes

### Competing Interest Statement

The authors have declared no competing interest.

### Summary of Updates

Figures have been modified to correct chromatic aberrations. The text has been edited accordingly.

