## Supplementary material for "SBIS, a new orange fluorescent vital probe for the 4D imaging of brown algal cells": Fig. S1

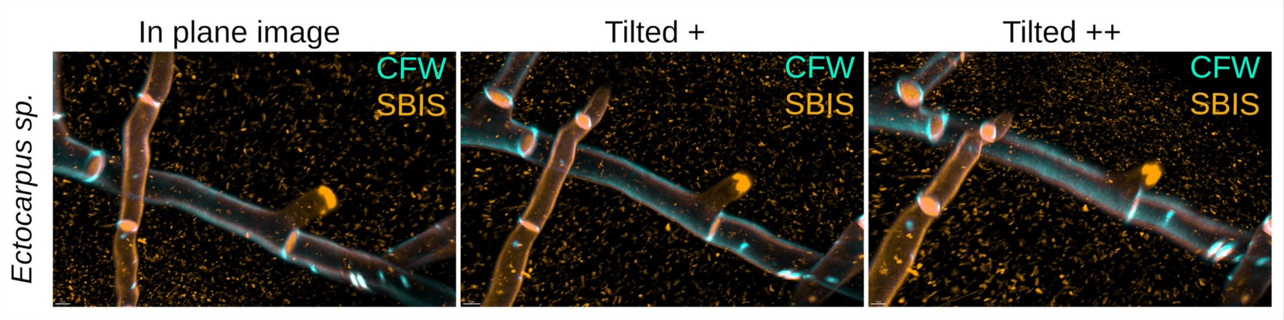


**Figure S1: Artifact in the observation of the location of SBIS in *Ectocarpus* filament cells.**

A 3D image of an *Ectocarpus* filament was reconstructed from a Z-stack acquisition using confocal microscopy, and then viewed using the Imaris 10.2.0 image viewing software (Bitplane, South Windsor, CT, USA). Depending on the sample orientation in the virtual reconstruction, the SBIS signal is located differently to the CFW signal. In the left image, SBIS is located internally compared to CFW in both longitudinal cell lines. This corresponds to an observation plane that is perpendicular to the median plane of the filament. Centre: in the bottom cell outline, SBIS is located externally compared with CFW. This corresponds to an observation plane that is slightly tilted compared to the perpendicular plane. Right: SBIS is located externally to CFW in both longitudinal cell outlines. This corresponds to a much more tilted observation plane compared with the perpendicular plane.
